# A Zebrafish Model for Ocular Tuberculosis

**DOI:** 10.1101/177576

**Authors:** Kevin Takaki, Lalita Ramakrishnan, Soumyava Basu

**Affiliations:** Molecular Immunity Unit, Department of Medicine, MRC Laboratory of Molecular Biology, University of Cambridge, Cambridge, CB2 OQH, UK; L V Prasad Eye Institute, Bhubaneswar, India

**Keywords:** tuberculosis, ocular, zebrafish, model, macrophage

## Abstract

Ocular tuberculosis (TB) commonly causes severe inflammation and vision loss in TB-endemic countries. The mechanism by which tuberculous infection becomes established in the eye is poorly understood. We used *Mycobacterium marinum*-infected zebrafish larvae to study the early pathogenesis of ocular TB and found hematogenous bacterial seeding of the eye despite a functional blood retinal barrier. Prototypical early granulomas formed that involved the retinal vasculature and retinal pigment epithelium-choroid complex; characteristic locations for human ocular TB. Peripheral blood monocytes were recruited to the growing granuloma suggesting that the immune privileged nature of the eye is breached by this inflammatory focus.

**Conflict of interest:** none disclosed

**Funding:** This work was supported in part by a ‘Short-term fellowship’ to SB by Department of Health Research, Government of India.

## INTRODUCTION

Ocular tuberculosis (TB) causes significant visual impairment in TB-endemic countries [1]. Several clinical presentations, involving nearly every ocular tissue, have been attributed to ocular TB [2]. The pathogenesis of ocular TB is poorly understood prompting several attempts to create animal models for investigation. These include rabbit models infected with *Mycobacterium tuberculosis* by intra-ocular or intra- carotid injections [3], and a recent model in aerosol-infected guinea pigs [4], both of which have demonstrated caseating as well as non-caseating granulomas in the eye, with acid-fast organisms. While these models have reproduced the histopathological changes of human ocular TB, the mechanisms by which mycobacteria invade the eye and how they establish infection in this immune privileged site (owing to the presence of the blood-retinal barrier) remains unknown.

The zebrafish (*Danio rerio*) model of *Mycobacterium marinum* infection has been established as a natural, relevant and genetically tractable host-pathogen conjugate, for dissecting mycobacterial interactions with the host in TB [5-6]. Zebrafish larvae are optically transparent, which allows real-time, intravital visualization of early host-pathogen interactions [5]. We reasoned that the zebrafish larva could be a useful model for studying the earliest host-pathogen interactions leading to ocular TB especially since the larval eye completes substantial anatomical development even by 2 days post-fertilization (dpf) [7]. The inner (endothelial) and outer (retinal pigment epithelium, RPE) blood retinal barriers (BRB) are well established by 3 dpf, rendering it a distinct anatomical compartment structurally analogous to the human eye [8]. Furthermore, the larva is highly amenable to intravital microscopy up to 3 weeks as well as to genetic manipulation [5].

## METHODS

### Fish husbandry

All zebrafish husbandry and experiments were performed in compliance with the UK Home Office and the Animal Ethics Committee of the University of Cambridge. Wild-type AB and various transgenic(Tg) zebrafish (*Tg*(*mpeg::YFP), Tg(mpeg:DsRed2) Tg(kdrl:DsRed2)* and *Tg(mfap4:tdTomato)* were maintained as described [6]. Eggs were collected following natural spawning and treated with N-Phenylthiourea 12 hours post-fertilization (hpf) onwards to prevent pigmentation.

### Bacterial strains

We used *M. marinum* strain M (ATCC BAA-535) constitutively expressing the green- or red-fluorescent proteins, Wasabi or tdTomato (pTEC15 or pTEC27 deposited with Addgene). *M. marinum* single-cell preparations for infections were made as described [6], and stored in 5 μL aliquots at a concentration of 100 CFU per nL. For each experiment, a single vial was thawed and mixed with 0.25 μL of 20% phenol red before injection.

### Caudal vein injections

Larvae at pre-determined developmental stages were anesthetized with tricaine and injected with *M. marinum*, or the specified reagent, as described [6].

### Microscopy

Screening of eyes for infection with fluorescent *M. marinum* was performed on a Nikon E600 equipped with a CoolLED pE-300 light source. Larvae were anaesthetised in tricaine and placed laterally on a flat slide to image one eye at a time. Each eye was imaged along the z-axis, from superficial to deep layers and the presence of fluorescent bacteria was scored manually. For confocal microscopy, larvae were embedded in low-melting point agarose (1.5%), and imaged on a Nikon A1 confocal microscope with a 20x Plan Apo 0.75 NA objective. Z-stack images were generated using a 2 µm step size spanning a total depth of 140 µm. Images were analysed using NIS-Elements AR 4.4.

### Peripheral monocyte recruitment assay

For these experiments, *Tg(mfap4:tdTomato)* fish with red-fluorescent macrophages were injected with a lower dose (25 CFU) of Wasabi-expressing green-fluorescent *M. marinum*. Fish were screened for granuloma formation at 6 dpi (9 dpf) and then injected with 200 mg/mL Hoecsht 33342 dye (Invitrogen). Recruitment of Hoechst-positive peripheral monocytes was evaluated by confocal microscopy 24 hours later.

## RESULTS

### Hematogenous dissemination of *M. marinum* infection to the eye

Since most forms of human extra-pulmonary TB are thought to develop through hematogenous dissemination of *M. tuberculosis* [9], we examined dissemination of *M. marinum* to the eye after intravenous infection of 3 dpf larvae (in which the BRB is already established). For this, we injected red-fluorescent *M. marinum* into the caudal vein of transgenic *mpeg:YFP* larvae with yellow fluorescent macrophages. We observed ocular infections in approximately 20% of the animals in multiple experiments, with the majority of infections already apparent 1 day post- injection (dpi) (table S1). Since an appreciable frequency of seeding to the eye was occurring in the presence of the BRB, it appeared that it was not a significant deterrent to bacterial traversal into the eye. To test this directly, we compared the frequency of infected eyes after 1 day of infection of 3 dpf larvae to 2 dpf where the BRB is not yet present [8]. If the BRB deters seeding of the eye, we should see an inhibition of ocular infection in the animals infected at 3 dpf. In contrast to our expectations, presence of the BRB did not inhibit bacterial seeding of the eye. In fact, the 3 dpf animals had a slightly higher frequency of infection (figure S1), possibly due to an increased number of bacteria traversing through the eye due to the increased retinal vasculature at this stage of development [8]. In any case, these findings showed that the BRB is not a significant deterrent to hematogenous seeding of mycobacteria into the eye.

### Kinetics of early granuloma formation in the eye

Sequential daily monitoring of the infection revealed the development of organized infected macrophage aggregates by 5 dpi (figure S2). The kinetics of retinal early granuloma development and their morphology were similar to that seen in other parts of the body [6,10]. In a few instances (∼5% eyes), we observed seeding of the eye at 1 dpi, with the infections not visible on repeat imaging, suggesting they had resolved or that the bacteria were transiently in the eye. In support of the former possibility, we have found that infection foci elsewhere in the body can also resolve, a phenomenon that has been observed in humans and nonhuman primates [11-12]. In support of the latter possibility, we noted occasional bacteria entering and leaving the eye through the hyaloid vasculature, suggesting that bacteria may transit through the eye without ultimately settling there. Overall, our findings suggest that mycobacterial infection of the eye results in the formation of macrophage aggregates with the same kinetics as elsewhere in the body.

### The anatomical localization of ocular granulomas in the zebrafish larva resembles that in humans

In humans, ocular TB is commonly localized to the retinal vasculature (retinal periphlebitis) or the retinal pigment epithelium - choroid complex (serpiginous-like choroiditis) (Figure 1 a-c). We found that mycobacterial infection was similarly localized in the zebrafish eye. Figure 1B showing the eye of a zebrafish with green transgenic macrophages infected with red-fluorescent bacteria, provides an example of an infected macrophage aggregate localized to the outer part of the eye, corresponding to the choroiditis lesion seen in the human eye. Figure 1C shows a *Tg(kdrl:DsRed2)* zebrafish eye with red-fluorescent retinal vasculature and green- fluorescent bacteria with bacteria abutting the blood vessels similar to retinal periphlebitis seen in human ocular TB. Since the macrophages in this animal are not labeled, we cannot tell if the bacteria are intracellular. It is likely that they are, given our observations in animals with labeled macrophages (figure 1B). In sum, the zebrafish lesions mechanistically represent the clinical phenotypes in human ocular TB.

**Figure 1.**
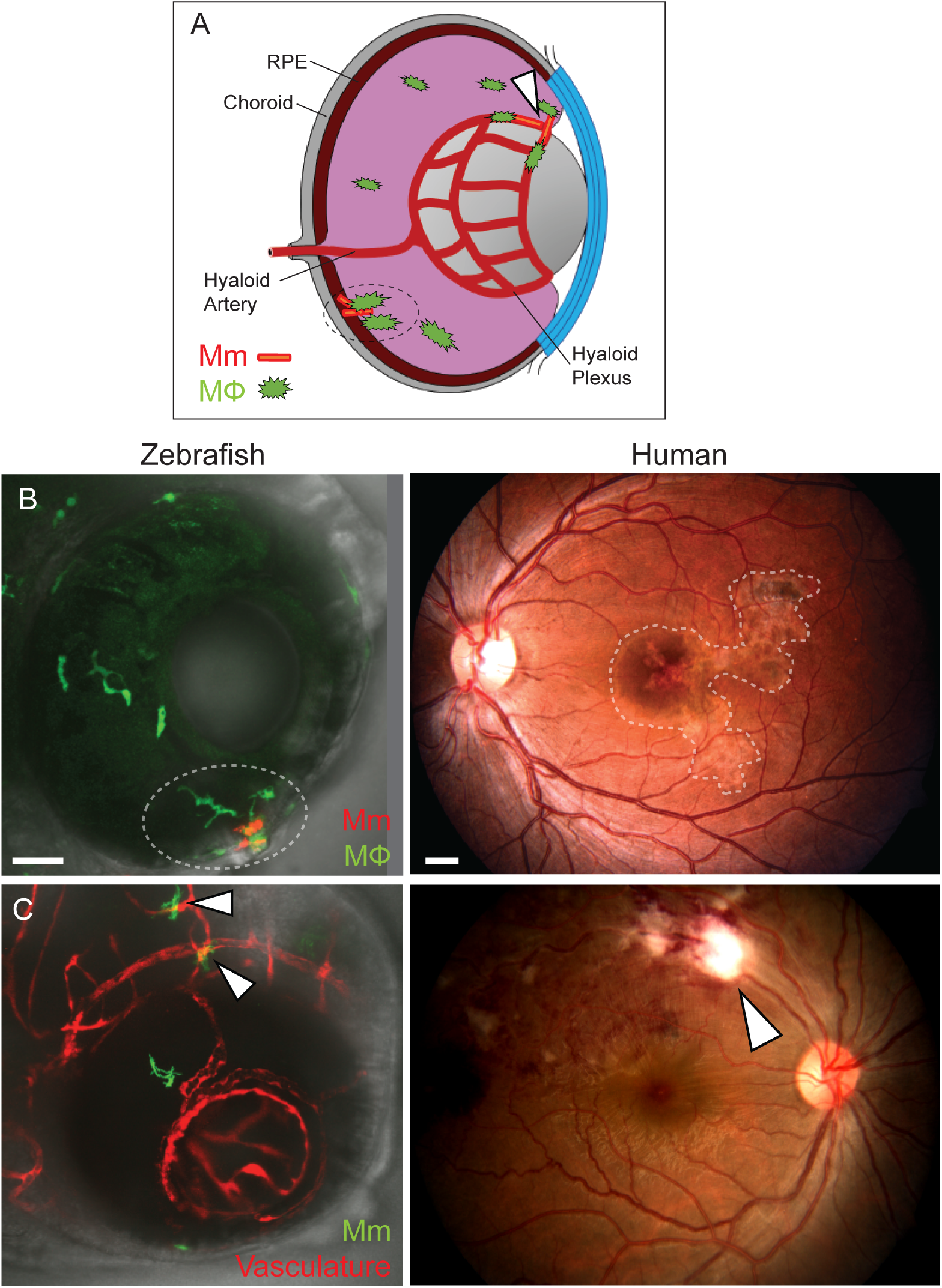
Anatomical localization of intraocular granuloma after *M. marinum* infection. (A) Schematic representation showing localization of granulomas near the retinal vasculature (arrowhead), and in retinal pigment epithelium-choroid complex (dotted circle). (B) Localization of intraocular granulomas (dotted circles); in the outer eye of zebrafish larvae corresponding to the retinal pigment epithelium-choroid complex, and in the choroid in human ocular TB. (C) Localization of perivascular infection (arrowheads); as seen as bacterial aggregates in close association of blood vessels in zebrafish, and retinal periphlebitis associated with focal chorioretinitis overlying the blood vessel in human ocular TB. (B-C) Confocal images of ocular infection in *Tg(mpeg:YFP)* and *Tg(kdrl:dsRed2)* zebrafish, and fundus photographs of human ocular TB. Scale bars, 25 μm and 750 μm, respectively.

### Ocular granulomas recruit peripheral monocytes

The eye has a population of tissue-resident macrophages (retinal microglia) that are already present even by this stage of development [13]. Their proximity to the site of infection would suggest that they participate in the formation of intraocular granulomas. However, we wanted to ask if peripheral blood monocytes were also recruited to the forming granuloma in the eye, as they are in the context of granulomas forming in other parts of the body [10]. To determine this, we took advantage of the nuclear dye Hoechst 33342 that on intravenous injection labels blood monocytes but not brain microglia owing to its inability to cross the BBB [10]. Next, we assessed zebrafish infected through caudal vein at 3 dpf for granuloma formation in the eye at 6 dpi, and injected the Hoechst dye in fish that had granulomas. Twenty-four hours after the injection, we found no dye leakage through the BRB, suggesting that there was not a gross breakdown of the BRB at this stage of the infection. When we imaged the developing granulomas, we found Hoechst- positive cells in 8 of 17 granulomas (47.1%) showing that they had recruited peripheral monocytes (figure 2A-C). Altogether, this experiment confirms that the BRB does not suffer gross breakdown even when sizable aggregates have formed. Whether there is localized breakdown of the BRB close to the lesions remains to be assessed as our imaging was not of sufficient resolution. Further, the findings that peripheral monocytes do not circulate into the uninfected eye are consistent with it being an immune privileged site. However, the relatively high frequency with which peripheral monocytes are attracted to the forming granuloma suggests that its chemotactic influence overrides the mechanisms that prevent monocyte ingress under baseline conditions.

**Figure 2.**
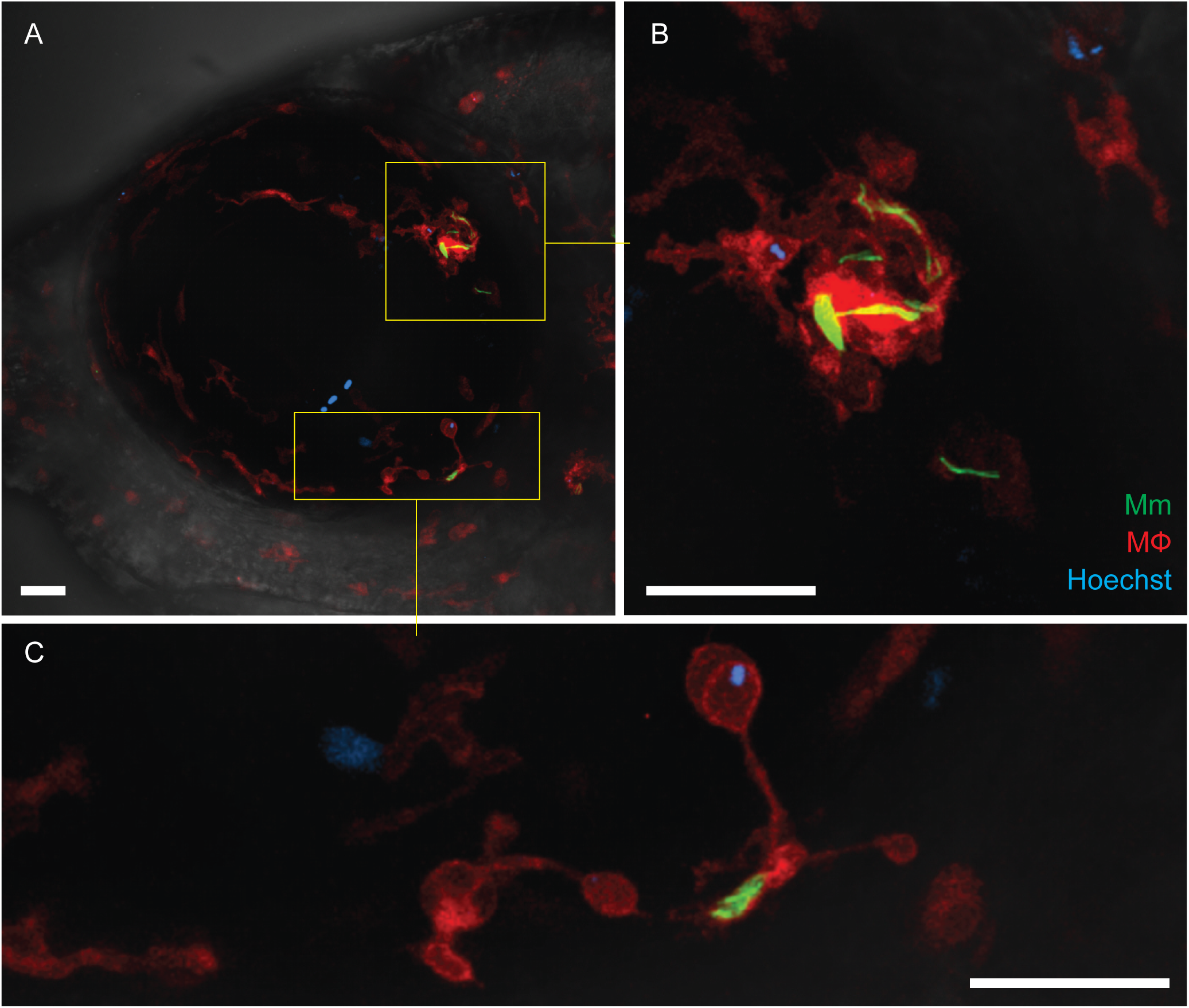
Recruitment of peripheral monocytes in granuloma formation. (A-C) Granuloma formation in *Tg(mfap4:tdTomato)* fish with red-fluorescent macrophages and infected with green-fluorescent *M. marinum*. Hoechst-positive (blue-fluorescent) circulating, non-tissue resident macrophages which have been recruited into the infected eye are seen within the granuloma (inset and panel B) and in contact with a single infected macrophage (inset and panel C). The remaining resident macrophages are dispersed uniformly within the ocular tissues. Scale bars, 30 μm.

### Discussion

We show that mycobacterial infection of zebrafish larvae can seed the eye hematogenously, consistent with the proposed model for extrapulmonary dissemination of TB in humans [9]. A large study of 10,524 cases of active pulmonary TB in a sanatorium, showed that 1.4% had ocular TB as well [14]. The much higher frequency of ocular dissemination (∼20%) in our experiments likely reflects the relatively high inoculum introduced directly in the blood stream. Alternatively, we may be looking at very early events that do not necessarily progress to clinically apparent lesions. In either case, the ability to study the first seeding event to the eye and the varying consequences thereof may shed light on these early events that have remained elusive so far. The similar spatial localization of human and zebrafish larval TB further validates the investigational utility of this model.

Two insights into the pathogenesis of ocular TB emerge from this pilot study. First, mycobacteria can enter the eye from the systemic circulation and establish infection despite the presence of an intact BRB. Hematogenous dissemination may explain the clinical finding that ∼ 50% of individuals presenting with ocular TB do not have evidence of TB elsewhere [2], suggesting that a transient bacteremia from an initial infection focus that resolved may have seeded the eye [9]. Second, once the infection is established in the eye, peripheral monocytes can be recruited from the circulation, despite limited or no breakdown of the BRB. We previously demonstrated the role of mycobacteria in intercellular expansion of infection through macrophage recruitment [10]. This work suggests that the monocyte-recruiting chemotactic signaling pathway induced by mycobacteria is operant in the face of the BRB, as it is in the context of the blood brain barrier. Resident macrophages, including brain microglia, are more microbicidal to mycobacteria than blood monocytes [15], and it will useful to see if this holds true in the context of retinal microglia and peripheral monocytes in the eye. It is conceivable that a preferential early involvement of the more proximate microglia in the forming ocular granuloma may tip the balance in the host’s favor to clear the infection.

We hope that this work will provide the foundation for future studies of the role of host and bacterial factors that influence dissemination to the eye and fate of the bacteria therein.

---

**Table S1:** Frequency of ocular dissemination (as % of infected eyes) after caudal vein injection of 100 CFU of *Mycobacterium marinum* in 3 days-post-fertilization zebrafish larvae, in three separate experiments.

**Figure S1. The blood retinal barrier does not prevent seeding of infection within eye.** Zebrafish larvae were infected with 100 CFU Mm at 2 dpf, before formation of the BRB, or at 3 dpf, following formation of the BRB. Each eye was scored for the presence of infection at 24 hours post-infection. Statistical significance determined by Fisher’s exact test.

**Figure S2:** Confocal image of green-fluorescent macrophage aggregate surrounding red-fluorescent *Mycobacterium marinum*, located in the retinal parenchyma. The mycobacteria are intracellular (inside macrophages) as well as extracellular (free). Scale bar, 50 μm.

